# LocusPackRat: a Semi-Automated Framework for Prioritizing Candidate Genes from Large GWAS Intervals

**DOI:** 10.64898/2025.12.03.692167

**Authors:** Brian Gural, Todd Kimball, Anh N. Luu, Christoph D. Rau

## Abstract

Genome-wide association studies (GWAS) routinely implicate broad loci that span tens of megabases and contain dozens of genes, making the leap from locus to causal gene challenging, especially in model organism cohorts with reduced mapping resolution. We developed LocusPackRat, a semi-automated, easily extendible package that assembles standardized ‘packets’ of evidence to accelerate candidate gene prioritization. Each packet merges study-specific information for each gene in a locus such as differential expression between conditions or presence of *cis-*eQTLs with functional/disease annotations pulled from InterMine and Open Targets. Packets are identically structured and easily disseminated to support side-by-side comparison and team review. We demonstrate LocusPackRat’s efficacy on a recent GWAS study of cardiac hypertrophy and failure in the Collaborative Cross. LocusPackRat shortens the path from statistical association to mechanistic hypotheses and improves the likelihood of successful experimental validation and is easily adaptable to other genetic reference populations or even human cohorts.

## Introduction

Genome-wide association studies (GWAS) and similar approaches (e.g. linkage studies) have led to the identification of many genes which are linked to phenotypes of interest ranging from height^1^ to blood pressure^2^ to cardiovascular disease risk^3^. A frequent bottleneck for these analyses is the transition from identifying loci to candidate genes. This is particularly the case in model organism genetic reference populations (GRPs) where mapping resolution is often lower and, consequently, loci routinely reach into the tens of megabases^4–7^ and contain dozens to hundreds of genes.

Identifying the best candidate gene from each locus often involves an effort to collate relevant information for each gene followed by locus-wide comparisons to prioritize the most likely candidate genes for downstream reporting and validation. As part of our work in the Collaborative Cross (CC), a murine GRP, we set out to create a semi-automated, easily extendable R package called LocusPackRat that standardizes and streamlines the aggregation of gene-level evidence to accelerate candidate gene nomination and validation. Below, we detail the components of LocusPackRat and apply it to a GWAS from the Collaborative Cross.

## Methods

Although these methods describe resources relevant to our application of LocusPackRat to the Collaborative Cross, equivalent resources exist for other GRPs.

### Desired qualities of high-confidence candidate genes (Figure 1B)

In a locus, there are often tens to hundreds of potential candidate genes to consider, any of which could, in theory, be one of the phenotype-driving genes at the site. Our goal, therefore, is to identify a handful (ideally one) of genes from each locus for further study. The ideal candidate gene would have the following characteristics:

**Figure 1.**
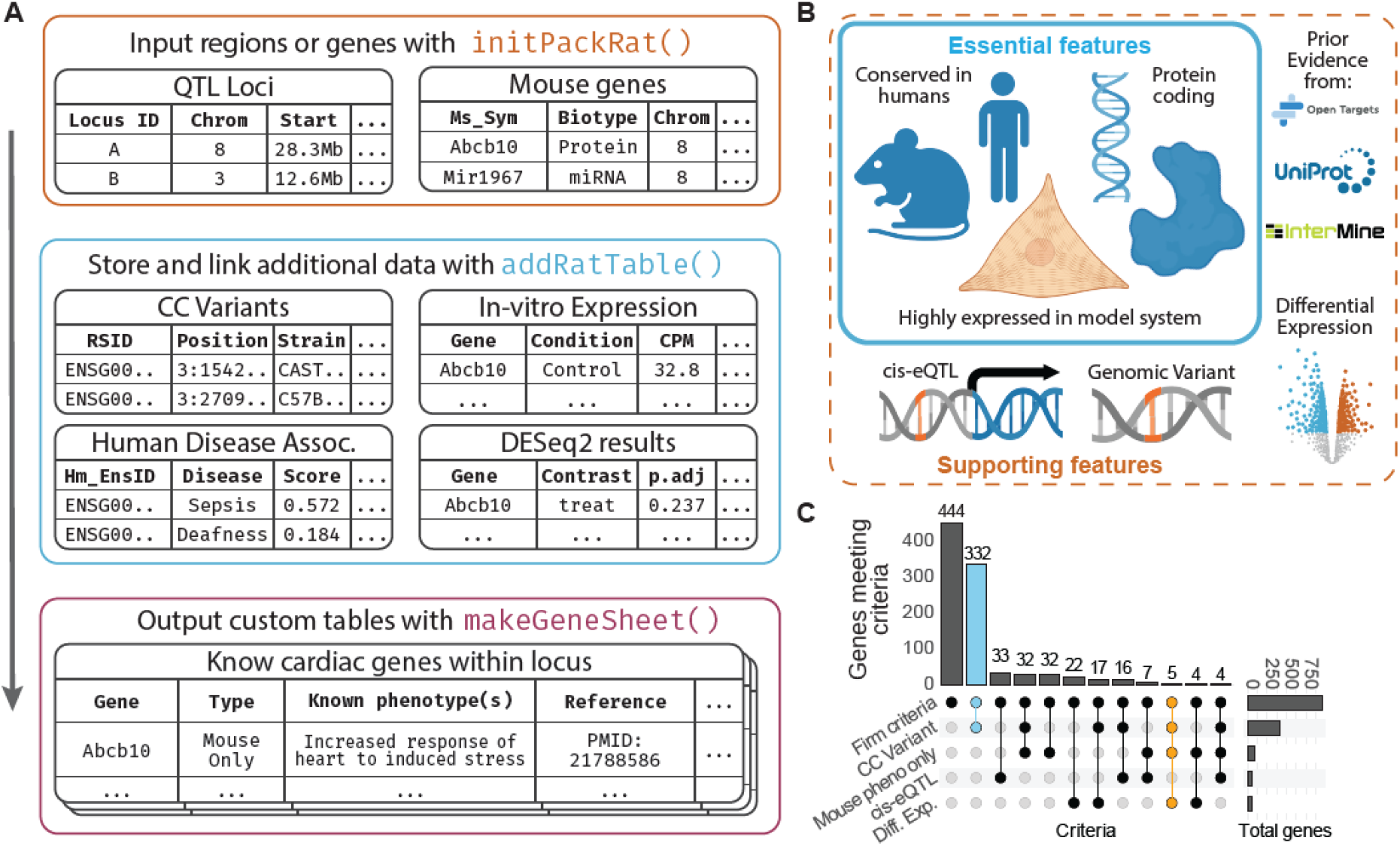
locusPackRat schema, application, and outcomes of evidence integration. **(A)** Users begin by supplying either QTL intervals or a gene list. initPackRat() creates a standardized “.locusPackRat” project (inputs, cache, supplementary, outputs), harmonizes identifiers (gene symbols/Ensembl IDs), and records genome/orthology metadata. Study- or resource-derived evidence is then attached with addRatTable(), which links supplementary tables to genes or regions. Finally, makeGeneSheet() takes inputs and user-provided criteria to generate “gene sheets” (preformatted CSV/Excel). **(B)** In a study of heart failure in mice, we considered essential features for candidate genes to include: presence of a human ortholog, protein-coding status, and robust expression in the intended model system. Supporting information was added from external sources and internal studies. **(C)** Example evidence overlap across 20 loci associated with heart failure phenotypes in mice. UpSet plot summarizing how many genes meet each criterion alone (right bars) and each combination of criteria (top bars). Connected dots beneath each bar indicate the included evidence types; “Firm criteria” corresponds to the essential features in (B). Three genes were selected from these results for downstream validation, Mrps5(blue), Lmod3, and Abcb10 (both yellow) in our full GWAS study

<u>Required:</u>

1. Expression in a relevant tissue/cell type
2. Experimentally actionable - protein-coding and with a human orthologue

<u>Supportive</u> (as many as possible):

1. Observed differential expression in response to a relevant stressor
2. Predicted deleterious non-synonymous variant
3. Detected significant *cis*-eQTL within the locus
4. Recovered prior links in literature or databases to mechanistically relevant traits in either an animal model or humans

### Structure of LocusPackRat (Figure 1A)

To assist in the identification of genes which meet these criteria, LocusPackRat creates ‘packets’ of information for each locus of a GWAS study, structured identically across all loci to enable direct comparisons, with standardized file naming and organization.

Packets include:

A. A multi-sheet Excel workbook containing
  a. A summary sheet providing basic information regarding the generation of the packet, formatting, and other sheet information
  b. A master sheet containing all genes within each locus that includes all information provided by the user (e.g. expression levels, differential expression, mutations, etc) along with information pulled from MouseMine^9^ and OpenTargets^10^ databases.
  c. Custom user sheets constructed by subsetting the master sheet
  d. Sheets from MouseMine and OpenTargets cleanly listing each linked phenotype, pubmedIDs of relevant literature, and the OpenTargets disease score
B. LocusZoom^8^-style visualizations displaying the GWAS locus and gene tracks, along with other relevant information, such as haplotype effect trajectories

### Mined Information

The mouse genetics community benefits from several large consortia committed to defining the function of every expressed gene in this model organism, with commitments by groups such as the International Mouse Phenotyping Consortium (IMPC)^13^ and the Knock Out Mouse Project (KOMP)^14^ to eventually create knockout models of each gene in the genome, although full completion remains a long-term goal. The Mouse Genome Informatics (MGI) website has compiled all current phenotype information into a searchable MouseMine database which runs off of the intermine framework and is queryable through the InterMineR package^15^. In a similar manner, up-to-date human disease associations have been consolidated in the Open Targets GraphQL API^10^. LocusPackRat returns all mouse and human gene-trait associations from these databases and has a customizable filter for specific traits of interest. We have found it useful to examine this information in two passes – first looking for genes explicitly tied to our phenotype of interest to look for known genes that may explain a locus, then a second pass to identify genes with functional evidence of roles in related phenotypes (e.g. high blood pressure vs cardiac hypertrophy) which may represent more novel or less well-studied candidates.

### Sequence-Level Information

The Wellcome Trust Mouse Genomes Resource^16^ provides basepair-level resolution for genetic variants in 36 mouse strains. These include the 8 founder lines of the CC, which, when paired with the known haplotype information for each CC strain allows us to identify specific point mutations in each CC line. Similar results could be achieved for many other mouse panels, such as the BxD cohort^6^ or Hybrid Mouse Diversity Panel^5^, each of which have their founder lines present in the sequencing data. Similar datasets exist for other GRPs^17^. As interpretation of non-coding mutations can be very difficult, we limited ourselves to non-synonymous, protein-modifying mutations for each protein-coding gene.

### Study-Specific Data

The data described above are of common interest to any study and can be used to filter candidate genes, however these lists can be further refined if researchers have additional, study-specific data in the form of phenotypic or transcriptional information, for example, ATAC-seq or CHiP-seq data for open chromatin regions or Hi-C linkages between the tag SNP and individual gene promoters, or simply transcriptional data from RNAseq or Microarray analyses.

For transcriptional data, we typically use two different analyses. First, differential expression between treated and control conditions (in our case Isoproterenol-treated(ISO) and saline-treated(Ctrl)) using DESeq2 to identify genes whose expression is significantly affected by our experimental system across all genetic backgrounds and highlights genes in loci that are significantly differentially expressed. Second, eQTL data to link individual gene expression to the locus.

## Results and Discussion

### Collaborative Cross Study (Figure 1C)

We now describe the application of LocusPackRat to our recently reported GWAS on isoproterenol(ISO)-driven cardiac hypertrophy and failure in the CC^11^.

Heart failure is characterized by a complex etiology in humans with multiple initial causes, but most of these inciting incidents eventually result in an increase in beta adrenergic signaling and catecholamine-driven overdrive of the heart which leads to failure^12^. We used the drug Isoproterenol, a synthetic beta adrenergic agonist, to induce heart failure phenotypes in 454 mice from 71 strains of the CC. We then mapped both control and ISO-treated phenotypes to the mm39 genome using the miQTL tool as described^11^. We identified 49 genome-wide significant loci, which consolidated to 20 distinct genomic intervals when overlapping regions and multi-trait loci were considered. These loci averaged 12.8Mb and contained a total of 2,149 genes, which we wanted to prioritize for experimental validation in neonatal rat ventricular cardiomyocytes (NRVMs). Below, we detail the results from the entire set of loci, as well as a specific locus on chromosome 3 which shows strong associations with cardiac echocardiographic traits.

### Initial Screening

We began by filtering our potential candidate gene pool based on the following criteria: 1) Genes must have obvious human orthologues (74% of all candidate genes), be robustly expressed in the cell type we intended to validate in (NRVMs, 46% of all candidate genes), and, lastly, be protein-coding to prioritize historically more easily manipulatable targets. These criteria gave us the best possibility of being able to query our candidate genes *in vitro* using *siRNA*-mediated knockdown or adenovirus-associated overexpression studies. This resulted in 950 total genes (44% of the original) that needed further curation.

### Manual Prioritization (Figure 1C)

After producing automated locus packets, the final step of candidate gene prioritization relies on manual curation by the researcher(s). Very few genes (only 1, in the case of the CC study) will possess evidence in all categories, while many genes will show evidence in a few. For example, we observed 407 genes that met our basic criteria and had a nonsynonymous variant in at least one founder strain, but only 50 which were differentially expressed between ISO and Ctrl cohorts. A final screen based on researcher domain expertise and with an eye for biological plausibility acts as a powerful final filter for gene validation.

We recommend approaching these prioritization tasks as a team effort. As LocusPackRat provides unified, consistent locus packets, we assigned two or three lab members (to minimize the effect of one individual’s personal biases towards specific pathways or degree of knowledge) to each locus, and each locus was examined for convergent evidence patterns, mechanistic plausibility, tractability for downstream testing in isolated cardiomyocytes, and translational potential for human cardiovascular studies. Results were presented in informal presentations in which evidence, both for and against each gene, was considered before a final prioritized gene candidate list was compiled for each locus.

### Examination of a Single CC GWAS Locus (Figure 2A,B)

As a concrete example, we apply LocusPackRat to a locus associated with post-ISO cardiac functional parameters on chromosome 3 that spans 19Mb (131-150Mb) and contains 104 genes. Of these, 50 pass our screening step (human orthologue, protein-coding, robustly expressed). Of these 50: 14 have a protein-altering mutation in PWK/PhJ, the most likely driver of this locus (mostly missense mutations but with 2 start_loss variants) while 7 other genes have protein-altering mutations in other strains, 10 have a *cis* eQTL at this locus, 5 show significant differential expression in response to ISO, and 18 have been previously linked to a cardiovascular-associated trait in MouseMine or OpenTargets. Only one gene (the beta-mannosidase enzyme Manba) had both a PWK-mutated protein-coding mutation *and* a cis-eQTL, while all but one differentially expressed gene had either a missense mutation or a *cis-*eQTL. 2 (40%) differentially expressed genes had been previously implicated in mouse or human for another cardiovascular trait, along with 6 (60% of total) *cis-*eQTL genes and 6 genes with non-synonymous mutations (29% of total). The entire packet may be found in the supplement.

**Figure 2.**
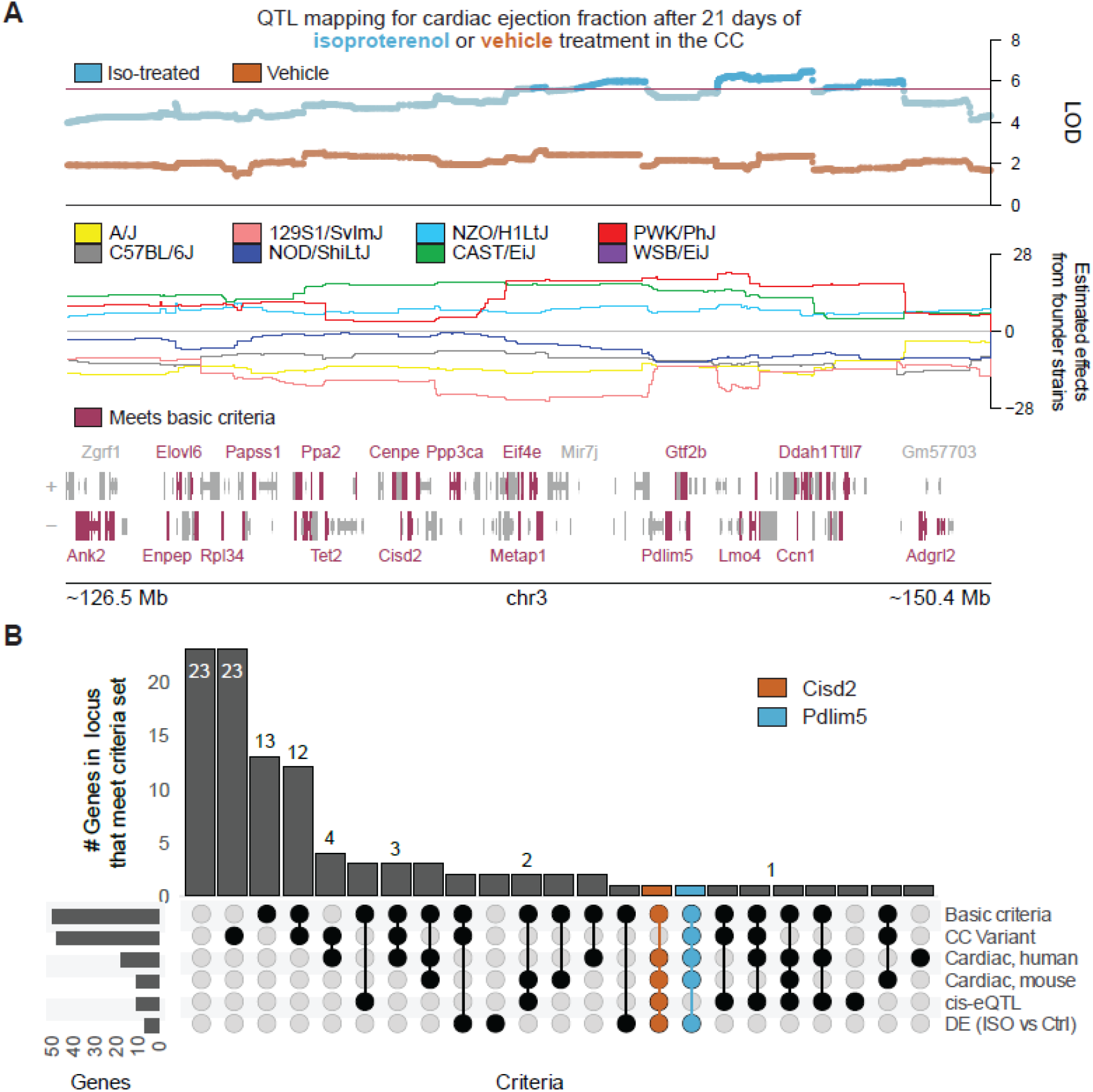
Example application of locusPackRat to a wide locus on mouse chromosome 3. **(A)** Top: Locus zoom plot showing the logarithm of the odds (LOD) for associations between genomic markers in the Collaborative Cross (CC) and measured cardiac ejection fraction after isoproterenol-induced stress (blue) or vehicle (orange). Dashed line indicates genome-wide significance threshold set by permutation testing. *Middle*: Estimated contributions of CC founder haplotypes to observed associations. *Bottom*: Gene tract showing genes within the locus (mm39). Genes that have a known human orthologue, encode protein, and are expressed above 5 cpm in NRVMs were prioritized for manual comparison with locusPackRat. **(B)** Upset plot showing the outcome of criteria set on all genes within the locus. Compelling candidate genes, Cisd2 and Pdlim5 are colored in orange and blue. CC Variant = Gene contained a coding variant in the CC; Cardiac, mouse = Reported association with cardiac traits in mouse models per Mouse Genome Informatics; Cardiac, human = Reported association with human cardiac phenotypes per Open Targets; cis-eQTL = detected as a locally regulated gene in internal eQTL mapping; DE: detected as differentially expressed between treated and control mice in internal analysis. Colored columns indicate columns which contain the two genes we chose to highlight from this cluster.

After considering the evidence, our attention settled on two genes as the most likely candidate genes for this locus. One is *Cisd2*, a regulator of autophagy that shows a strong *cis-*eQTL at this locus, differential expression between control and ISO conditions, and has been implicated in abnormal skeletal muscle morphology, although its role in cardiac muscle remains to be explored^18^. The other is *Pdlim5*, a scaffolding protein whose overexpression may lead to dilated cardiomyopathy^19^ and which has a nonsynonymous mutation in PWK/PhJ in the CC and significantly increased expression after ISO. Next steps would involve examining each of these genes in an *in vitro* system for further validation.

### Summary and Code Availability

LocusPackRat is a semi-automated R package meant to facilitate the often complicated steps between the identification of a genome-wide significant locus and the identification of the key gene or genes that underly the locus. By providing a standardized ‘packet’ of information that consolidates information from the original study, species-or-cohort-specific data such as sequencing data, and gene-level information such as known functions and knockout effects, LocusPackRat allows for teams of researchers to work together to prioritize candidate gene lists with everyone having the same data at their disposal, speeding the process of picking candidates.

LocusPackRat is implemented as a modifiable and extendable R package at github.com/RauLabUNC/locusPackRat.

### Limitations

We have extensively tested LocusPackRat in the context of the analysis of heart-related CC-derived loci, and its efficacy in other cohorts may be limited by different sources of information available. LocusPackRat was designed to be easily extendable to other GRPs by simple modifications to incorporate panel-and-phenotype-specific information. For maximal efficacy, researchers should have user-supplied sequence-level DNA information and RNA transcriptomes from their study. For many organisms (flies, humans, etc), equivalent resources to MouseMine exists through intermine.org (e.g. FlyMine). Each missing data source will likely lead to a reduction in the utility of LocusPackRat, and an increased reliance on user intuition and longer research times to identify the best candidate genes. Furthermore, the murine community has greatly benefitted from the efforts of groups like the IMPC or KOMP which have performed preliminary phenotypic screens of many knockout lines. Other model organisms with fewer resources or more limited gene annotations will further find the efficacy of LocusPackRat limited, yet it should still provide a foundation for further efforts.

## Abbreviations

CC: Collaborative Cross
DE: Differentially Expressed
(e)QTL: (expression) Quantitative Trait Locus
GRP: Genetic Reference Population
GWAS: Genome-Wide Association Study
IMPC: International Mouse Phenotyping Coalition
ISO: Isoproterenol
KOMP: Knock Out Mouse Project
MGI: Mouse Genome Informatics
NRVM: Neonatal Rat Ventricular Cardiomyocyte
SNP: Single Nucleotide Polymorphism

## Declarations

### Ethics Approval

Not Applicable

### Consent for Publication

Not Applicable

### Availability

Phenotypes for the CC may be found in the Mendeley Data Repository at the following location: data.mendeley.com/datasets/vyf5x4ygrv/1

LocusPackRat is available at Github: github.com/RauLabUNC/locusPackRat

### Competing Interests

The authors declare they have no competing interests

### Funding

This work was supported by R01HL162636 and R00HL138301 (CDR) and T32HL069768 (BG)

### Author Contributions

**BG** conceived of the LocusPackRat tool, wrote the package, conducted the analyses and interpretation of the data, and drafted the manuscript

**TK** contributed to the acquisition of the CC data

**AL** contributed to the analysis of the CC data

**CDR** oversaw the research, contributed to the acquisition and interpretation of data, and drafted the manuscript

All authors contributed to revisions

## Acknowledgements

Not Applicable

